# Complementing Muscle Regeneration: Fibro-Adipogenic Progenitor and Macrophage-Mediated Repair of Elderly Human Skeletal Muscle

**DOI:** 10.1101/2024.04.18.590053

**Authors:** Jonas Brorson, Lin Lin, Jakob Wang, Tine Borum Billeskov, Jesper Just, János Haskó, Christen Ravn, Rehne L. Hansen, Mats Bue, Yonglun Luo, Niels Jessen, Jean Farup

## Abstract

The capacity to regenerate skeletal muscle function after injury requires a complex and well-coordinated cellular response. Here, we unravel the intricate dynamics of human skeletal muscle regeneration by combining spatial, temporal, and single cell transcriptomics. Using spatial RNA sequencing, we profiled the expression of human protein-coding genes in elderly human skeletal muscle biopsies before as well as 2-, 8-, and 30-days post injury. Single Cell-Spatial deconvolution analysis highlights monocytes/macrophages and fibro-adipogenic progenitors (FAPs) as pivotal players in human muscle regeneration. By utilizing flow cytometry and cell sorting we confirmed increased cellular content and activity during regeneration. Spatial correlation analysis unveils FAPs and monocytes/macrophages co-localization and intercellular communication, mediated by complement factor C3. Immunostaining confirms C3 expression in FAPs and FAP secretion of C3, suggesting a role in phagocytosis. Functional assays demonstrate C3’s impact on monocyte metabolism, survival and phagocytosis, unveiling its involvement in skeletal muscle regeneration. These insights elucidate FAP-macrophage interplay with perspectives for future therapeutic interventions.

## Introduction

Skeletal muscle represents the predominant tissue in the human body, constituting up to 30-50% of total body mass in healthy, lean individuals (1). Its functions encompass pivotal roles in breathing, locomotion, thermoregulation, and glucose homeostasis, while also serving as a modulator of immune activity (2). Thus, the preservation of skeletal muscle quantity and quality across the lifespan is indispensable for maintaining human health. Skeletal muscle exhibits remarkable adaptability and, when faced with acute injury, demonstrates remarkable regenerative capacity. Even severe and repeated injuries in healthy young muscle results in complete restoration of muscle morphology and function (3, 4). However, this regenerative process is curbed during disease (5), and impaired ability to regenerate skeletal muscle mass is a central feature of the aging process.

Restoration of muscle tissue integrity and function post-injury relies on the intricate coordination between multiple cell types to facilitate phagocytosis and clearance of necrotic muscle cells, scaffold generation, stem cell expansion, and differentiation, culminating in the replacement of damaged parenchyma without excessive collagen deposition (6, 7). During the initial phases after injury, the recruitment of leukocytes to the injury site is crucial for the initiation of normal muscle regeneration (8). Chemokines, complement factors, and cell adhesion molecules play pivotal roles in attracting leukocytes following muscle injury (9–12). For instance, depletion of Chemokine Ligand 16 decreases macrophage infiltration, impairs regeneration resulting in fibro-fatty replacement of muscle tissue (13). In a physiological response to injury, pro-inflammatory macrophages secrete TNF-alpha which induces apoptosis of fibro-adipogenic progenitors (FAPs) (14), and this prevents excessive FAP proliferation and collagen deposition. In contrast, anti-inflammatory macrophages secrete TGFβ, which prevent FAP apoptosis and promotes collagen production (14) and if unchecked, may ultimately impair regeneration and lead to fibrotic replacement of muscle tissue. Collectively, there is compelling evidence of leukocyte regulation of FAP proliferation and differentiation during muscle regeneration.

Conversely, several lines of evidence suggest that FAPs may play a crucial role in recruiting macrophages to the site of injury. In mice, genetic depletion of FAPs severely reduces regenerative capacity (15, 16). Although the mechanisms underlying this phenomenon remain elusive, it has been reported that FAP-secreted IL6 and WNT1-inducible-signaling pathway protein 1 (WISP-1) (17, 18) support expansion and myogenic commitment of muscle stem cells. Additionally, genetic depletion of FAPs leads to a marked impairment in the clearance of necrotic muscle cells and reduced influx of immune cells (16), significantly impeding muscle regeneration. Evidence from non-muscle tissues, such as the synovial membrane, suggests that subsets of fibroblasts can indeed initiate repeated synovial inflammatory processes (19). However, while leukocytes in skeletal muscle are evidently involved in controlling both expansion and differentiation of FAPs after injury, direct evidence supporting the reverse is currently lacking.

Insights into the cellular response to injury is predominantly based on rodent models, but some observations of an interplay between leukocytes and FAPs in human skeletal muscle have been reported. It is reported that fibroblasts proliferate following injury in human skeletal muscle (20), and when muscle regenerate, infiltration of macrophages is observed in close proximity to regenerating muscle fibers and proliferating muscle stem cells (21–24). This could suggest that FAPs and macrophages communicate directly during regeneration of human skeletal muscle.

In this investigation, we illuminate the intricate cellular and inter-cellular dynamics of human skeletal muscle regeneration through the integration of spatial and single-cell transcriptome analyses with the aim to identify putative signaling mechanisms of cellular communication. Using an experimental design with repetitive biopsies from healthy elderly subjects, we are able to describe the chronological order of cellular events during physiological conditions and unravel a novel FAP-macrophage communication.

## Results

### Spatial and temporal transcriptome analysis of human skeletal muscle regeneration

We used a unique human model of electrically induced eccentric muscle contractions of the vastus lateralis part of the quadriceps femoris muscle (3, 25) to obtain substantial tissue injury and subsequently regeneration in aged subjects (**Figure 1A**). As muscle regeneration occurs in highly coordinated and time-dependent manner, skeletal muscle biopsies were obtained before (pre) as well as 2, 8, and 30 days post injury (dpi) to uncover different phases of the regenerating process. H&E-staining revealed disrupted tissue integrity, infiltrated myofibers, and increased cell content at 8 dpi (**Figure 1A**). Tissue integrity was predominantly restored at 30 dpi although myofibers were smaller, containing more central nuclei, and expressing embryonic myosin heavy chain (shown previously (25)). Collectively, we confirm myofiber injury, removal, and subsequent fiber regeneration in this model of human skeletal muscle injury.

**Figure 1:**
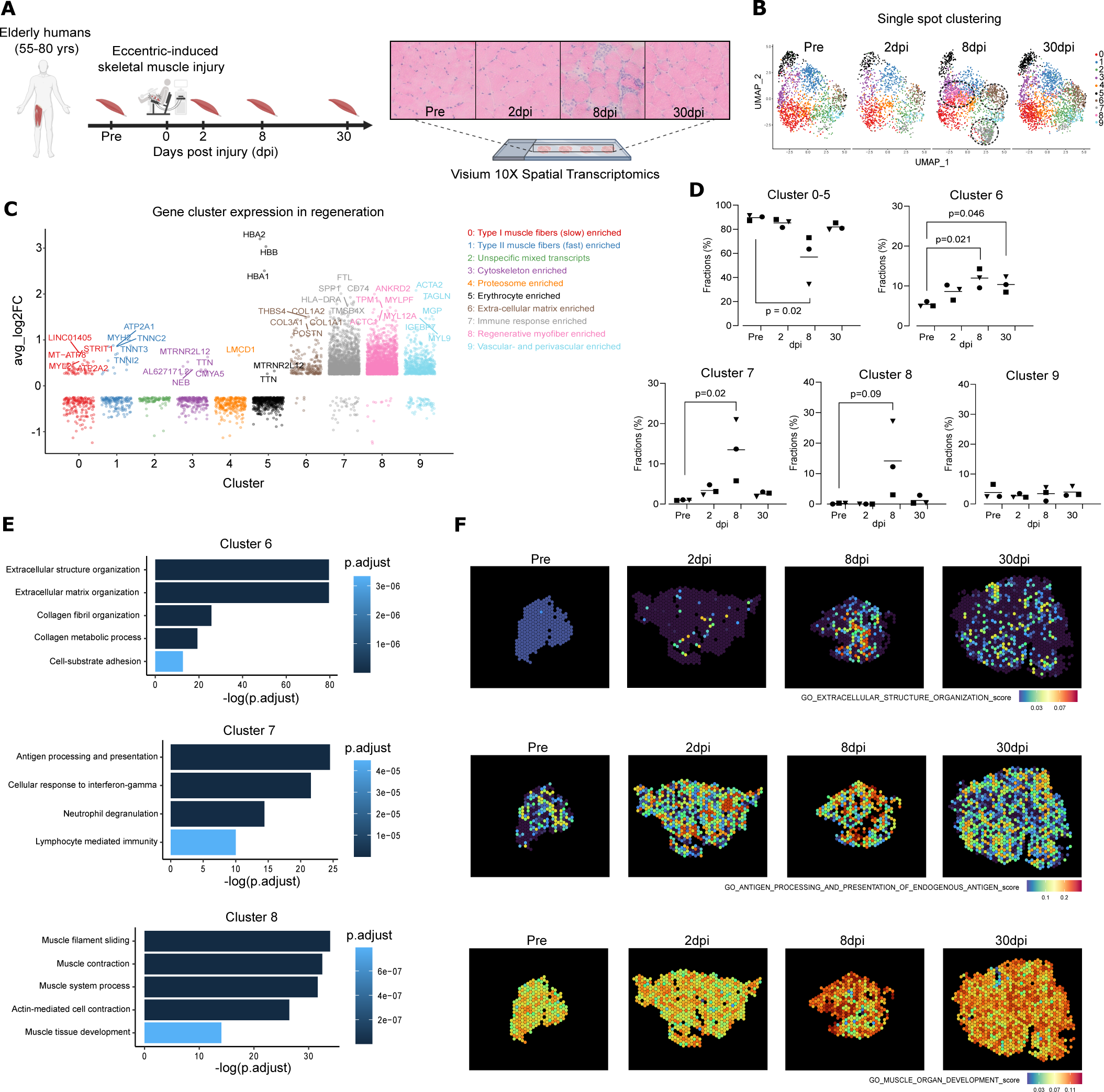
Spatial transcriptome analysis of regenerating human skeletal muscle. A: Schematic illustration of the muscle injury protocol and analysis. Elderly human subjects were recruited and exposed to electrically induced skeletal muscle injury in the vastus lateralis part of the quadriceps femoris muscle wherefrom biopsies were obtained before (pre), 2, 8, and 30 days post injury. H&E staining of cryosections confirmed substantial tissue injury. Spatial transcriptome analysis was carried out in muscle tissue from three individual subjects using the 10x Visium platform. B: Clustering of single spatial tissue spots during the time course of regeneration. Spots were independently divided in 9 clusters based on expression pattern. Every dot represents a single spatial spot. Spots containing transcripts from cluster 6, 7, and 8 were highly present after injury (marked). C: Change in gene expression during regeneration based on gene clusters. Expression in cluster 0-5 was decreased whereas cluster 6, 7, and 8 r evealed an increase in gene expression after injury. D: Fraction of total spots constituting cluster 6 and 7 were increased after injury and cluster 8 tended to be increased. E: Gene Ontology analysis of cluster 6, 7, and 8 shows enrichment of genes related to extracellular matrix, immune response, and myofiber development, respectively. F: Representative spatial expression of top gene clusters from Gene Ontology analysis with wide spatial increase most intensive 8 days post injury. dpi = days post injury.

To gain deeper insight into spatial and temporal gene expression changes during muscle regeneration, we used the Spatial Gene Expression technology (10x Genomics Visium) to profile all human protein-coding genes expression in the skeletal muscle biopsies. The Visium Spatial Gene Expression technology is based on spatially barcoded oligonucleotides that capture messenger RNA (mRNA) molecules from the partially permeabilized tissue sections (26). We first optimized the permeabilization protocol to achieve homogeneous mRNA release without disrupting the histological structure of muscle tissues. cDNA staining showed that 24 min permeabilization of human skeletal muscle (10 µm thickness) can homogenously release the mRNA from the tissue without compromising its morphological structure (**Supplementary Figure 1A**). We then performed spatial gene expression profiling of the muscle biopsies samples at pre, 2, 8, and 30 dpi from three individuals. The mean reads per captured spot under the tissue were 93,828. After filtering spatial plots with low Unique Molecular Identifier (UMI) count (total UMI < 700), we obtained in average 620, 412, 757, and 761 spatial spots from pre, 2 dpi, 8 dpi, and 30 dpi, respectively. Notably, both median genes and median UMI detected per spatial plot were significantly higher at 8 dpi than other time points (**Supplementary Figure 1B**), corroborating the dynamic cellular environment in this stage of regeneration.

We first sought to explore the spatial expression pattern in muscle injury and regeneration. Unsupervised clustering was performed with the merged top 3000 most highly variable genes (HVG) within each sample and the top 30 principal components. Using Uniform Manifold Approximation and Projection (UMAP)-based dimension reduction and visualization, we identified ten spatial clusters with distinct gene expression patterns (**Figure 1B**). Based on differentially expressed genes, geneset enrichment analyses and spatial projection (**Supplementary Figure 2A**), we annotated the spatial clusters according to their enriched biological functions. Two spatial clusters were identified as muscle fibers: Type 1 muscle fibers (slow, cluster 0) and Type 2 muscle fibers (fast, cluster 1), which were decreased (8 dpi) after muscle injury and restored at 30 dpi. Due to the presence of unspecific mixed transcripts, cluster 2 was not functionally annotated. Cluster 3 was enriched in cytoskeleton functions, which was decreased after injury. The functional cytoskeleton of muscle cells is vital for muscle structure, organization, and movements. In agreement with the muscle injury model, cluster 4 was enriched in proteasome functions and increased at 2 dpi, representing the extensive remodeling of the injured muscle cells. Cluster 5 was enriched in transcripts related to erythrocyte functions, illustrated by the top five expressed genes (**Figure 1C**). Collectively, we defined all these six spatial clusters (0–5) as injury-related clusters, which exhibit a pattern and functions related to injury (cluster 4) and injury-induced muscle remodeling (cluster 0-3, 5) and was significantly reduced at 8 dpi (Figure 1D, p = 0.02, one-way ANOVA). In contrast, cluster 6, 7, and 8 showed marked increase over the regenerative time-course, whereas cluster 9 (enriched in vascular- and perivascular transcripts, (**Figure 1C**) appeared relatively static (**Figure 1D**). The dynamic clusters 6, 7, and 8 were enriched in extracellular matrix organization (e.g. *COL1A2, COL3A1*, and *POSTN*), immune response (e.g. *CD14, CD163,* and *HLA-DRA*), and regenerative myofiber transcripts (e.g*. DES*, *MYH3*, and *NES*), respectively, and displayed a large temporal increase with all reaching a maximum 8 dpi (**Figure 1D**). We focused on the three spatial clusters (6, 7, and 8) exhibiting strong correlation with muscle regeneration and conducted Gene Ontology analysis and further validations.

Cluster 6 was highly enriched in extracellular structure and organization (**Figure 1E**), which corroborates the continued elevated ECM formation at 30 dpi (**Figure 1D**, p = 0.02 (8 dpi), p = 0.046 (30 dpi) as compared to pre), as extracellular matrix remodeling is still active in the late phase of regeneration (27, 28). The spatial distribution of genes in extracellular structure organization showed homogenous distribution in the early (2 dpi) and late (30 dpi) phase (**Figure 1F**). In contrast, at 8 dpi, where the largest gene expression changes were observed, there was marked clustering within few, spatially confined areas, which appeared to be interconnected (**Figure 1F**). Cluster 7 was highly enriched in genes involved in immune cell functions, such as antigen processing and presentation as well as interferon-gamma signaling pathways (**Figure 1E**). The cluster revealed a slight increase in the early and late phase of regeneration (2 dpi and 30 dpi) but with a marked increase at 8 dpi (**Figure 1D**, p = 0.02 compared to pre). Similar to cluster 6, the expression in early and late regeneration was less restricted to specific areas (although more than cluster 6), however, at 8 dpi there was strikingly local and spatial confinement of the changes in cluster 7. Importantly, the cluster 8 exhibits enriched functions and processes related to myofiber generation/regeneration, such as muscle system processes and muscle development (**Figure 1E**). In support of this, the expression pattern of this cluster corresponds to a tightly regulated process which is only, but intensely, increased at 8 dpi (**Figure 1D**, p = 0.094 compared to pre). While there was also some level of spatial restriction in the expression pattern at 8 dpi, this appeared to be less pronounced compared to cluster 6 and 7. Overall, these findings confirm a substantial regenerative response with different temporal and spatially confined gene expressions in the present injury model. To facilitate future studies and enhance our understanding of human muscle regeneration, we have generated a publicly available database (https://dream.au.dk/tools-and-resources/humdb) based on ShinyCell (29) for interactively exploring the spatial expression of protein-coding genes in this unique elderly human muscle regeneration model.

### FAPs and macrophages demonstrate spatial and temporal association in regeneration

To examine the spatiotemporal response to muscle injury of specific mononuclear cells, we deconvoluted all 7,650 spatial transcriptome spots based on a previously published single-cell RNA sequencing data from human skeletal muscle (30) using SPOTlight (31) (**Figure 2A**). This enables us, for the first time, to examine changes within all major subsets of mononuclear cells during regeneration of human skeletal muscle. In general, the deconvolution analysis revealed a striking coherence between our data and the typical cellular patterns observed in chemically induced muscle injury in mouse models (32, 33). Analysis of spatial spots containing cell-specific gene transcripts showed that monocyte/macrophage transcripts exert the largest response to injury, reaching a maximum at 2 dpi (**Figure 2B**). This is in agreement with the finding that monocyte/macrophage is the predominant leukocyte in single-cell RNA sequencing data from unperturbed human skeletal muscle (30, 34–36) and in early chemically induced injury in mice (33). Muscle stem cell (MuSC) transcripts were also rapidly increased at 2 dpi (**Figure 2B**), indicating an early activation of MuSCs. This finding further supports the enhanced MuSC cell cycle entry that we recently reported (37). Similar to macrophages and MuSCs, vascular endothelial cell-specific transcripts were also rapidly increased at 2 dpi. Unlike macrophages, MuSCs and vascular endothelial cells, the levels of FAPs, lymphocyte, and lymphatic endothelial cell transcripts reached a maximum at 8 dpi (**Figure 2B**). This reveals a highly coordinated and orchestrated cell activation during muscle injury and regeneration.

**Figure 2:**
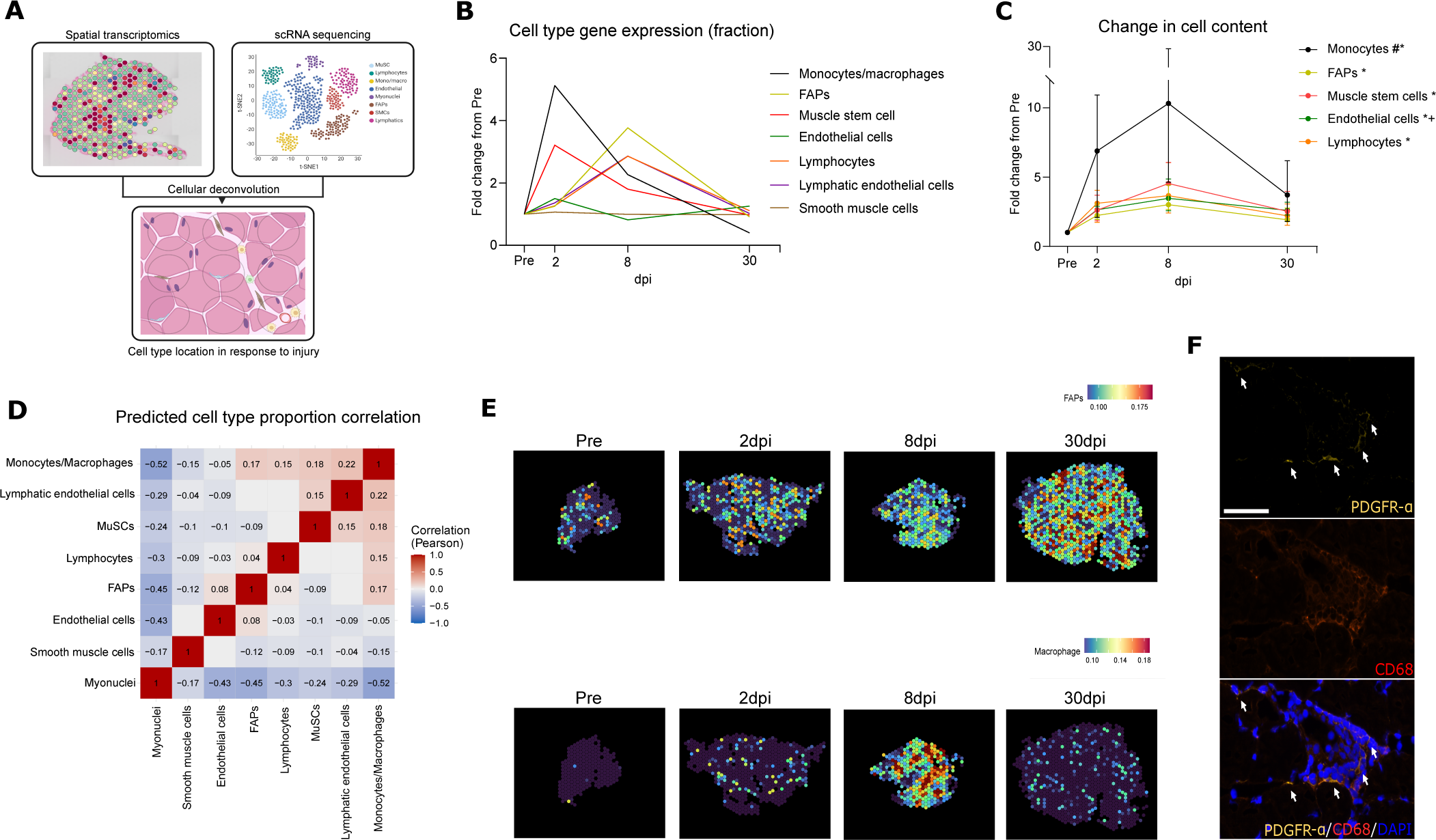
Cellular deconvolution and spatiotemporal associations in regeneration. A: Schematic illustration of the cellular deconvolution approach using sp atial transcriptomic and single -cell sequencing data from human skeletal muscle. B: Fractional distribution of specific cell type transcripts during the course of regeneration shows FAPs and monocytes/macrophages to constitute the two cell populations with largest relative increase(n = 3, curves represents means). C: Change in cell content during regeneration quantified from tissue homogenate by flow -cytometry. Monocyte content was increased in the early phase of regeneration whereas lymphocyte, MuSC, FAP and endothelial cell content was increased at later time points (# = p < 0.05 pre vs. 2dpi; (* = p < 0.05 pre vs. 8dpi; + = p < 0.05 pre vs. 30dpi) (Monocyte, MuSCs, endothelial cells, lymphocytes: n = 10; FAPs: n = 9) (graph represents median ± interquartile range). Cell-cell proportion correlation revealed that FAPs were spatiotemporally correlated with monocytes/macrophages, lymphocytes, and endothelial cells (D) (n = 3). E: Spatial distribution of spots containing FAP and monocyte/macrophage

To determine if the predictions of cellular content based on spatial and single-cell transcriptome analysis translate into cellular content, we quantified monocytes, FAPs, MuSCs, endothelial cells, and lymphocytes, in whole tissue homogenate by flow-cytometry. Monocyte content was increased at 2 dpi and further increased to 18-fold above baseline at 8 dpi (**Figure 2C**). In effect, the monocyte population constituted approximately 40% of total hematopoietic cells at 8 dpi compared to only 15% in uninjured conditions (**Supplementary Figure 3A**). This confirms a role for monocytes/macrophages also in human skeletal muscle regeneration. The relatively large and rapid increase in monocyte content likely reflects a major infiltration from circulating monocytes in agreement with earlier reports (8). FAP, MuSC, endothelial cell, and lymphocyte content were increased at 8 dpi (as well as 30 dpi for endothelial cells) (**Figure 2C**). Thus, the temporal cellular content showed a general alignment with the temporal activation of cellular gene expression obtained by the decomposition analysis.

To study the colocalization of cells during the muscle regeneration progress, we performed colocalization analysis by calculating the correlation (Pearson) of deconvoluted cell fractions between each two cell types within the spatial spots. We performed colocalization analysis for all eight major cell types across the 7,650 spatial transcriptome spots. Given our interest in FAPs as a sensor of tissue homeostasis, we focused cell types positively correlating with the localization of FAPs. Interestingly, our unbiased correlation matrix clearly indicated co-localization between FAPs and monocytes/macrophages (**Figure 2D**). While FAPs were positively correlated (Pearson r = 0.17) to monocytes/macrophages, monocytes/macrophages also exhibited high colocalization (Pearson r = 0.15 to 0.22) with other cell types including MuSCs and lymphocytes (**Figure 2D**). Given that macrophages and FAPs were two cell populations with the greatest spatial response to injury and were spatiotemporally correlated in regeneration, we further characterized a potential cellular interaction between these cell populations. Firstly, from our spatial RNA sequencing data, we noted a somewhat similar temporal response of FAP and macrophage spot transcripts but also with some spatial heterogeneity (**Figure 2E**). Second, to support our spatial transcriptome data, we performed immunostaining of PDGFR-α (FAP marker) and CD68 (macrophage marker) and found that cells positive for these markers reside in a close proximity in injured areas (**Figure 2F**). Interestingly, a similar spatial pattern of FAPs (PDGFR-α) and macrophages (CD68) has also been observed in degenerating skeletal muscle in patients suffering from Duchene’s muscular dystrophy (38). Overall, these findings demonstrate a close spatiotemporal relationship between FAPs and monocytes/macrophages in human skeletal muscle regeneration.

### Complement factor C3 is a key mediator for FAP to macrophage communication

To investigate cellular interaction and communication patterns between FAPs and monocytes/macrophages, we performed receptor-ligand analysis between muscle cell types based on human skeletal muscle single-cell RNA sequencing data (30). This revealed a large number of interactions from FAPs to endothelial cells, lymphatic endothelial cells, and monocytes/macrophages (**Figure 3A**). When exploring the details of these ligand-receptor pairs, we observed a diverse array of signaling between FAPs and monocytes/macrophages. Here the *C3* to *ITGAX+ITGB2*/*ITGAM+ITGB2* appeared as specific receptor-ligand pairs in the interaction between FAPs and monocytes/macrophages (**Figure 3B**). Not only was this pathway strongly enriched in the FAP-monocyte/macrophage communication, the C3 signaling also appeared specific for communication between FAPs and monocytes/macrophages (outgoing signal from FAPs and incoming to monocytes/macrophages) (**Figure 3C**). C3 is a central part of both the classical, lectin, and alternative complement pathway and whole-body ablation has been shown to perturb the early phase of monocyte/macrophage infiltration and skeletal muscle regeneration in animals (12). The hepatocytes are thought to be the primary source of circulating complement factors (39) although recent studies have reported that local C3 production plays an essential role in tissue inflammation and remodeling (40, 41). To investigate FAPs as a potential cellular origin of local C3 in human skeletal muscle, we first quantified the expression of *C3* in the different muscle cell types. Here, *C3* gene expression was almost entirely restricted to the FAP cluster (**Figure 3D**). To confirm the *C3* expression at protein level in human FAPs as well as C3 secretion, we isolated and plated FAPs from human skeletal muscle by FACS using established protocols (30, 36, 42). Immunostaining of C3 revealed that freshly isolated human FAPs were positive for C3 protein (**Figure 3E**). To further confirm that human FAPs secrete C3 in the microenvironment we collected conditioned media from freshly isolated FAPs, MuSC, and endothelial cells. Analysis of the conditioned media from FAPs revealed an ability to secrete C3 immediately after isolation which was strikingly lowered 48-96h after isolation, indicating a tight regulation of this property (**Figure 3F**). Transcriptome data also confirmed that local *C3* expression was increased early after injury (2dpi and 8dpi) (**Figure 3G**). In sum, these data show that FAPs from human skeletal muscle can both produce and secrete C3, however, the C3 expression and secretion is highly dependent of the state of the FAPs. At 48h-96h hours post isolation, the human FAPs are engaged in the cell cycle (30), suggesting that C3 is downregulated in the process of cell cycle.

**Figure 3:**
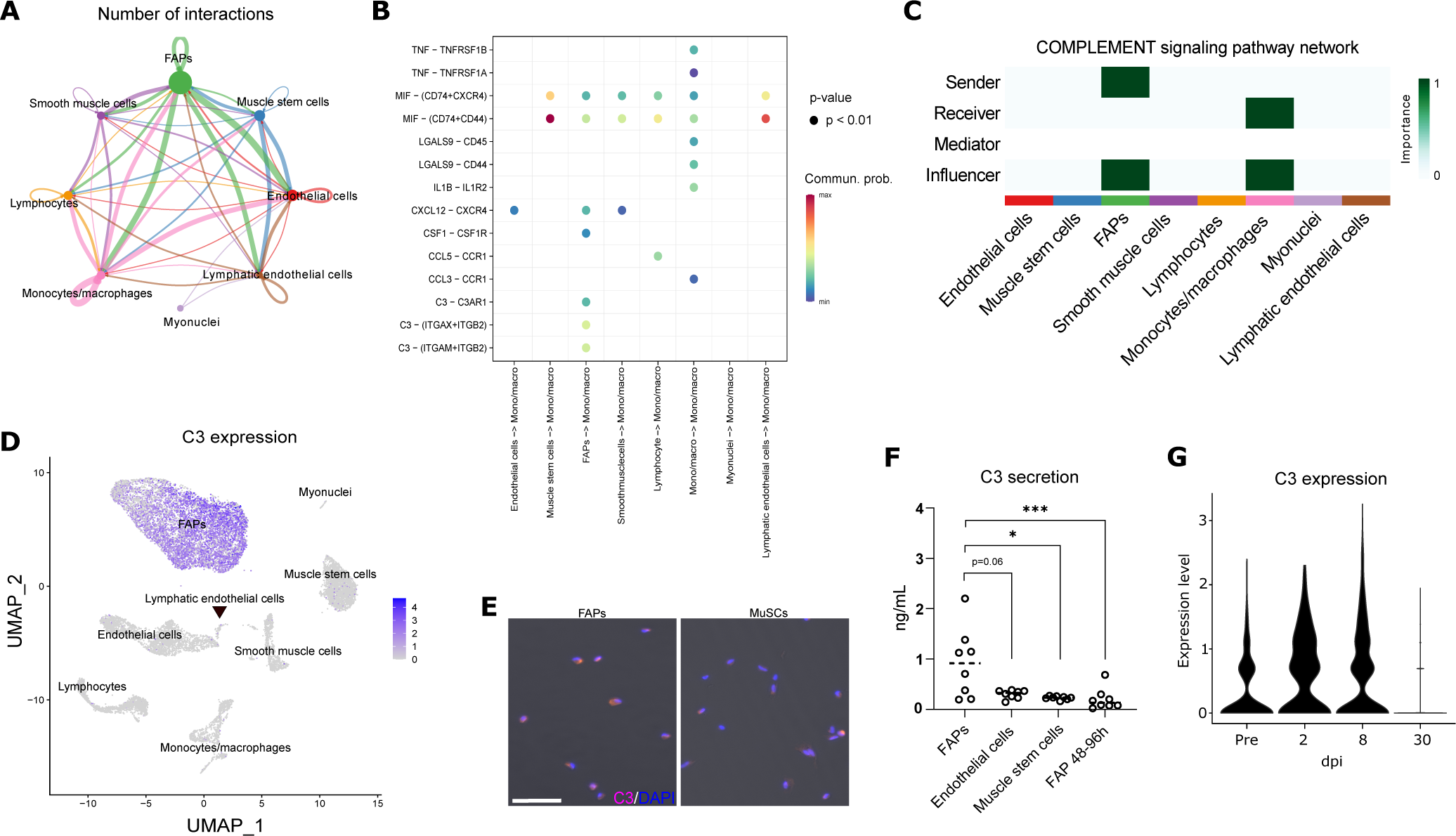
FAP derived comp lement factor C3 is a key mediator for monocyte/macrophage interaction. A: Receptor-ligand analysis of mononuclear cells from human skeletal muscle identifies diverse communicative pathways from FAPs to endothelial cells, lymphatic endothelial cells, and monocytes/macrophages. B: Communicative pathways from resident mononuclear cells to macrophages shows a large degree of interaction between FAPs and monocytes/macrophages with C3 - and MIF -pathways having the highest probability. C: Computed centrality scores heatmap of complement signaling in human skeletal muscle is relatively peculiar f or FAP -monocyte/macrophage communication, which is also shown by FAP expression of C3 compared to other mononuclear cell populations (D). E+F: FAP ability for C3 production and secretion was confirmed by C3 immunostaining and ELISA of C3 in conditioned media

The incoming signal from C3 to monocytes/macrophages was predicted to be mediated through *ITGAX+ITGB2* (and *ITGAM+ITGB2*) as revealed in our receptor-ligand analysis. *ITGAX* (CD11c) constitutes a subunit of the complement receptor 4 (together with *ITGB2*/CD18) which has C3 fragments as natural occurring ligands and are expressed on macrophages (43). Interestingly, we found that the majority of macrophages in the injured muscle were positive for CD11c protein (**Supplementary Figure 4A**) and flow-cytometry data confirmed a profound increase in CD11c^+^-monocytes starting at 2dpi and then further increasing at 8 dpi (**Figure 4A, B**). In effect, the ratio of CD11c^+/-^-monocytes increased from approximately 1 (before injury) to 3 (2 dpi) (**Supplementary Figure 4B**). Interestingly, while the CD11c^+^-monocyte content was increased at early time points, the CD11c^-^-monocyte content did not increase until 8 dpi and remained elevated at 30 dpi, which suggests differentiated roles in regeneration. Supporting this, we observed *ITGAX* expression was able to polarize the monocyte/macrophage population in human skeletal muscle (**Supplementary Figure 4C**), in an independent data set. Finally, this aligns with recent findings reported from our group that CD11c^+^-monocytes from human skeletal muscle are highly pro-inflammatory compared to CD11c^-^-monocytes based on cytokine secretion (36).

**Figure 4:**
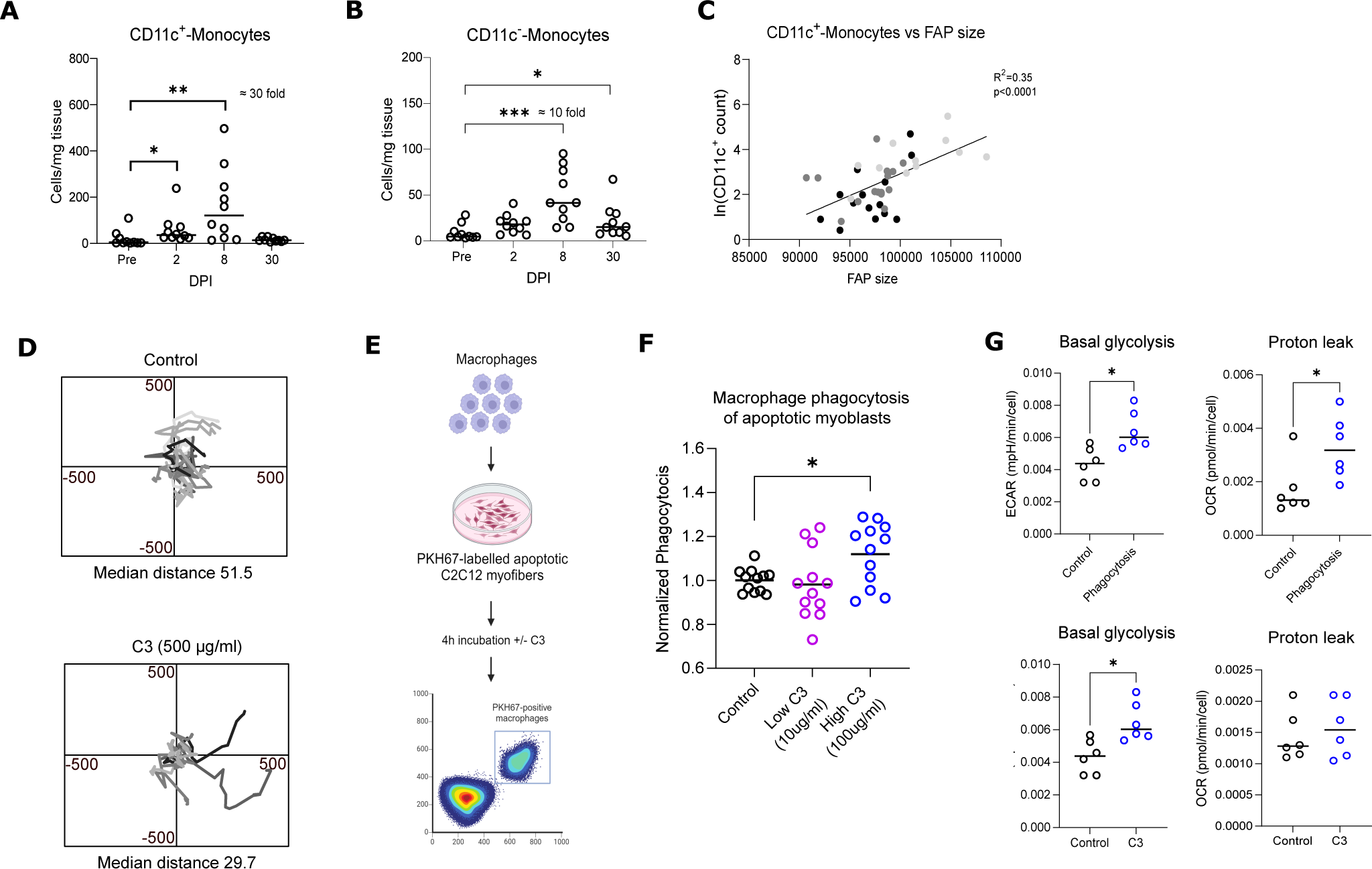
Complement factor C3 and associated monocytes/macrophages in muscle regeneration. A+B: Tissue content of CD11c+-Monocytes were increased in the early phase of regeneration whereas CD11c — Monocytes content were increased later in regeneration (n = 10). * = p < 0.05. ** = p < 0.01. *** = p < 0.001C. : Association between FAP size and CD11c+ -monocyte count indicates that tissue content of pro -inflammatory monocytes partly depends on FAP size. D: C3 p rotein decrease migration of monocytes. E: Schematic i llustration of macrophage phagocytosis experiment using apoptotic fluorescent labeled C2C12 myofibers and flow cytometry. Macrophage phagocytosis of apoptotic myoblasts showed that presence of C3 protein increased phagocytic activity. G: Macrophage

In unperturbed homeostatic muscle, FAPs are mainly assumed to be quiescent and only activated followed by cell cycle entry upon muscle injury or growth factor stimulation (30, 44). This activation increases cell size followed by DNA-synthesis and ultimately mitosis in a process that mirrors MuSC activation (45). In the present model of regeneration, we found FAP cell size increased 2 and 8 dpi and EdU-incorporation tended to be increased 8 dpi (**Supplementary Figure 4D+E**). This confirms that human FAPs are indeed activated by muscle injury. If FAP-derived C3 serves as an opsonization agent for macrophages in skeletal muscle regeneration, we hypothesized that FAP activation would associate with CD11c^+^-monocyte content. We used the early increase in FAP cell size as a marker of FAP activation and found that this parameter was highly correlated to CD11c^+^-monocyte content in skeletal muscle (**Figure 4C**). In support of this finding, Wosczyna and colleagues showed that ablation of FAPs decreased the early infiltration of hematopoietic cells (3 dpi) whereas later time points were unaffected (16).

Next, we asked which role C3 could play in human skeletal muscle regeneration. First, we were able to detect a close spatial proximity of C3 together with FAPs and macrophages in injured areas by immunostaining (**Supplementary Figure 5A**) which could indicate a direct role of C3 on macrophages. Our immunostaining also revealed injured myofibers positive for C3 protein. These myofibers were surrounded by macrophages suggesting a role in repair/phagocytosis of the injured myofiber (**Supplementary Figure 5B**). Following secretion, C3 protein is often cleaved by enzyme complexes (convertases) to form C3a and C3b (46). C3a exerts chemotactic properties as shown by decreased macrophage trafficking after injury in C3aR ablated animals (12). Conversely, C3b often appear surface-bound to form a C3/C5-covertase (46) and can act as an effective opsonization agent to target pathogens or apoptotic/necrotic cells for phagocytosis. The C3 antibody utilized in our staining detects the N-terminal part of C3 (C3b after cleavage). To investigate if the detected C3 was bound to the cell membrane of myofibers, we performed a co-staining with complement factor Bb. This demonstrated that factor Bb was highly abundant in injured muscle (**Supplementary Figure 5C**), which indicates opsonization of injured myofibers with C3b in the alternative complement pathway. To gain insight into the role of C3 in recruitment and later phagocytosis in human muscle injury we directly isolated human monocytes for either immidiate experiments or stimulated these into macrophages (CD68^+^) (**Supplementary Figure 6A+B**). The presence of C3 reduced monocyte migration/movement (**Figure 4D + Supplementary video 1**) which was somewhat in contrast to our initial expectations. However, we reasoned that reduced migration could be related to an increased opsonization and phagocytosis of dead cells and cellular debris. This speculation was supported by a markedly different monocyte morphology when stimulated with C3 in low serum conditions (larger and more spindle-shaped cells) (**Supplementary Figure 6C**). To test this hypothesis, we utilized a lipophilic dye to label C2C12 myoblasts and induced these towards programmed cell death (apoptosis). The apoptotic C2C12’s was then added to macrophages stimulated with low or high concentrations of C3 (**Figure 4E**). While low C3 did not alter the phagocytic activity we did observe a significant increase with high C3 levels (**Figure 4F**). Notably, the high C3 was still below normal plasma levels. In addition, we observed that C3 promoted survival of monocytes during a cellular stress situation stimulated through serum starvation (**Supplementary Figure 6C+D**). Combined with the effect of C3 on cell morphology and phagocytosis, this suggest a mechanism of C3 to support monocyte survival and phagocytosis through monocyte metabolism (47). To address this hypothesis, we generated macrophages from monocytes and shortly exposed these to apoptotic C2C12 myoblasts before evaluating glycolytic flux and oxygen consumption. Indeed, phagocytosis did alter macrophage metabolism with increased glycolytic activity as well as marked increased proton leak (**Figure 4G**). An increased uncoupling (proton leak), likely mediated through UCP2, has previously been demonstrated to be critical in supporting effective phagocytosis in macrophages (48). Interestingly, C3 alone, without apoptotic C2C12 myoblasts, also effectively stimulated a shift towards increased glycolysis in macrophages (**Figure 4G**) with no significant increase in proton leak and maximal oxygen consumption, although we did observe a significant increase in respiratory reserve capacity (**Supplementary Figure 6E**). This metabolic switch is an analogue to C3 meditated metabolic reprogramming of synovial fibroblasts which is important in sustained tissue inflammation (40). Overall, these findings support a direct role for C3 in phagocytosis and metabolism (12, 40).

Since the majority of our understanding of skeletal muscle regeneration and the mechanistic C3 investigations have been performed in mice, we sought to investigate if C3 was also specifically expressed in murine FAPs. We explored specific FAP *C3* expression in mice skeletal muscle before and after injury at a single-cell level based on data published by De Micheli, Laurilliard (33). These confirmed a specific expression of *C3* in FAPs before injury (**Supplementary Figure 7A+B+C**). After injury, FAP specific *C3* expression was increased in the early phase of regeneration and returned to baseline levels 7dpi (**Supplementary Figure 7C**). Expression was also increased in innate leukocytes after injury, and this might contribute to C3 signaling although FAPs constitute the quantitative largest reservoir for C3 also in murine skeletal muscle.

Collectively, our findings support a model in which FAPs act as local source of C3 to initiate presence of pro-inflammatory monocytes/macrophages in injured human skeletal muscle supporting local phagocytosis. These findings support the essential need for properly functioning FAPs to mount a successive regenerative response in skeletal muscle.

## Discussion

Using a combination of single-cell and spatial transcriptomics on muscle biopsies sampled before and several times after injury, we have uncovered a novel intercellular communication axis between FAPs and macrophages during human skeletal muscle regeneration. We identify FAP-derived complement factor 3 as a key signaling molecule capable of attracting and supporting the phagocytotic function of pro-inflammatory monocytes/macrophages. The identification of essential intercellular communication in tissue remodeling offers the opportunity to target these cells and intercellular pathways for supporting human skeletal muscle in ageing and disease associated with loss of muscle mass, function, and regenerative capacity.

The ability of FAPs to serve as a local origin for C3 production and secretion within human skeletal muscle is notably intriguing. While the absolute level of C3 expression and C3 secretion in FAPs is likely lower than the levels secreted from hepatocytes (55?) this does not preclude functional importance, particular in local tissue injury. Notable, in this study, we did not observe any changes in systemic (plasma) levels of C3 during the course of muscle injury, suggesting a more localized role for C3 production in FAPs. C3 expression has been determined in FAPs or FAP-like cells (mesenchymal stem cell, stromal cells, fibroblasts) in a variety of organs including cardiac, kidney and adipose tissue (49, 50). Comparisons to data from mice suggest that this mechanism is conserved among other vertebrates indicating that the local production serve a fundamental biological function to aid and control tissue inflammation and regeneration. While the clinical implications for altered C3 regulation is yet unclear, it has been demonstrated that age-related loss of muscle mass, also termed sarcopenia, is associated reduced plasma levels of C3 in elderly subjects (51). The association between C3 and the mass and function of human skeletal muscle could suggest that either FAPs, FAP derived C3 or both, are involved in the development of sarcopenia in elderly subjects. Moreover, given the critical function of the innate immune system in muscle regeneration, this also implicate reduced C3 in the impaired regenerative capacity associated with aging or inactivity. In mouse skeletal muscle, the complement system, in particular C3, is of key importance for skeletal muscle regeneration and homeostasis (12, 52, 53). It is demonstrated by constitutive whole-body ablation or inhibition of C3, as well as deletion of complement factor B and C3a-receptor, that C3 is necessary for macrophage chemoattraction and thus successful regeneration of injured muscle (12). Similarly, the essential role of FAPs in successful muscle regeneration is well-established in mouse models (15, 16), although the underlying mechanism has remained insufficiently understood. However, the delayed clearance of necrotic muscle fibers combined with the markedly reduced inflammatory infiltrate, indeed indicated a link between FAPs and chemotaxis or function of immune cells (16). We found that C3/C3b positive human myofibers were surrounded by macrophages after injury which indicates a role for C3 in opsonization and debris removal, as suggested in animal models (12). Moreover, our in vitro data support a role for C3 in phagocytosis of apoptotic muscle cells, collectively proposing that the complement system is essential in both monocyte recruitment and clearance of damage muscle cells, also in humans. Dysregulation of this intercellular communication in pathology is so far speculative, although the association between sarcopenia and C3 does indicate clinical consequences for muscle mass and regenerative capacity (51). Conversely, in chronic cycles of muscle regeneration/degeneration during neuromuscular pathologies such as Dysferlinopathies (53) depletion of C3 provide beneficial restoration of muscle function. Considering the close association with accumulation of FAPs in degenerative skeletal muscle disorders, this supports a close association between local muscle inflammation and FAPs. Furthermore, recent evidence has implicated C3 secretion from synovial fibroblasts as a key mechanism underlying tissue priming and recurrent inflammation in affected joints in rheumatoid arthritis (40). Notably, the latter also reported a metabolic priming of synovial fibroblasts mediated by C3, with a marked increase in aerobic glycolysis of the C3 primed fibroblasts. Similarly, we also observed a generalized metabolic priming of macrophages by C3, with a particular increase in aerobic glycolysis. Interestingly, we found similar alterations in the context of phagocytosis, suggesting that the metabolic changes could support phagocytotic capacity of the macrophages. Besides macrophages, FAP have also been reported to be endowed with phagocytotic capabilities of necrotic muscle cells (54), suggesting that the effect of C3 may not be restricted to monocytes/macrophages. Collectively, the accumulating evidence indicate that pathological regulation of local C3 expression can predispose to chronic local tissue inflammation.

A recent study by Babaeijandaghi et al. revealed direct communication between FAPs and macrophages in skeletal muscle through Colony stimulating factor 1 (CSF1) (55). They report, that a specific subset of FAPs expressing DPP4, secrete CSF1 to maintaining a pool of self-renewing macrophages in mouse skeletal muscle. By investigating our transcriptomics data, we observed that CSF1 and C3 seems to be expressed in different subsets of FAPs (https://dream.au.dk/tools-and-resources/humdb) supporting that they are likely required for different condition, i.e. maintaining homeostasis or supporting the regenerating muscle niche. Thus, FAPs are likely involved in regulating inflammation in both homeostatic conditions and in acute muscle injury.

Understanding human skeletal muscle regeneration has posed challenges due to limitations in generating robust muscle injury models. For instance, the nature of human skeletal muscle regeneration is often characterized by more focal areas containing necrotic myofibers while other areas are morphologically unaffected (4, 22). Our study capitalizes on spatial transcriptome analysis to map transcripts specifically from regenerating areas, providing unprecedented insights into human muscle regeneration. By integrating spatial transcriptome spots with single-cell RNA sequencing data, we have delineated a role for FAPs in macrophage recruitment during human skeletal muscle regeneration. Validation of these findings through flow cytometry analysis further strengthens the robustness of our approach, offering a unique perspective on human muscle regeneration with implications for future hypothesis generation and target validation.

In conclusion, our investigation employing single-cell and spatial transcriptomics techniques has revealed significant alterations in FAP and macrophage content, along with close spatial and temporal associations between them during human skeletal muscle regeneration. We have identified several FAP-secreted ligands, including C3, which interact with pro-inflammatory monocytes/macrophages, suggesting their pivotal role in facilitating optimal skeletal muscle regeneration through mechanisms of monocyte recruitment, opsonization, and debris removal.

## ACKNOWLEDGEMENTS

Flow cytometry/cell sorting was performed at the FACS Core Facility, Aarhus University, Denmark. Illustrations were created using Biorender.com. This study was funded by grants from the A.P. Møller Foundation given to JB, A.P. Møller Foundation, Riisfort Foundation, Toyota Foundation, The Independent Research Fund Denmark (DFF – 5053-00195) given to JF and the Steno Diabetes Center Aarhus, which is partially funded by an unrestricted donation from the Novo Nordisk Foundation. We acknowledge the Novo Nordisk Foundation (NNF17OC0027242 to NJ).

## AUTHORS’ CONTRIBUTIONS

Conception and design of research: JB, LL, YL, NJ, and JF. Performed experiments: JB, JF, JW, TBB, CR, RLH, MHB, and LL. Analyzed data: JB, JJ, LL, YL, and JF. Interpreted results of experiments: JB, LL, JJ, JW, YL, NJ, and JF. Prepared figures: JB, LL, JJ, YL, and JF. Drafted manuscript: JB, LL, YL, NJ, and JF. Approved final version of manuscript: All authors.

## DECLARATION OF INTEREST

The authors declare no competing interests.

## Methods

### Ethical approval

The study was conducted in accordance with the Declaration of Helsinki after approval by the local Research Ethics Committee in Region Midtjylland, Denmark (1-10-72-301-18 and 1-10-72-308-20). The muscle injury study was registered at clinicaltrials.gov (NCT03754842) before recruitment was commenced. All participants received oral and written information before written consent was obtained.

### Study population and sample preparation

Analysis of skeletal muscle injury and subsequently regeneration was carried out on skeletal muscle samples from the placebo group of a previously published study (25). In brief, skeletal muscle injury was induced in a dynamometer by 200 involuntary muscle contractions initiated by electrical stimulation of the vastus lateralis part of the quadriceps femoris muscle simultaneously with a greater opposite force applied by the dynamometer arm (eccentric muscle work). Skeletal muscle biopsies were obtained from the vastus lateralis part of the quadriceps femoris muscle with manual suction under local analgesics (Xylocain 10 mg/ml, AstraZeneca, Stockholm, Sweden) in sterile conditions. A minimum distance of 3 cm was kept between each incision to minimize the effect of repeated sampling. After removal, parts for histology were aligned and embedded in Tissue-Tek® O.C.T compound (Sakura Finetek USA, Inc.) and frozen in precooled isopentane followed by transportation in liquid nitrogen before storage at −80°C. Part of the biopsy for FACS, were immediately put into a C-tube (cat. no.: 130-093-237, Miltenyi Biotec, Lund, Sweden) containing ice cold wash buffer (HAMS F10 incl. glutamine and bicarbonate (cat. no.: N6908, Sigma-Aldrich, Denmark) + 1% Penicillin-Streptomycin (cat. no.: 15140122, Gibco, ThermoFisher Scientific, MA, USA) + 10 % horse serum (cat. no.: 26050088, Gibco, ThermoFisher Scientific, MA, USA). The C-tube was weighted before and after the biopsy was added to obtain total milligram of skeletal muscle. Before digestion, Collagenase II (cat. no.: 46D16552, Worthington, Lakewood, NJ, USA) and Dispase II (cat. no.: 04 942 078 001, Roche Diagnostics, Basel, Switzerland) were added to the C-tube making a final concentration of 700 U/ml and 3.27 U/ml, respectively. The biopsy was then digested using a gentleMACS (cat. no.: 130-096-427, Miltenyi Biotec) for 60 min at a skeletal muscle program (37C_mr_SMDK1). Afterwards, 10 µl of wash buffer was added before filtering through a 70 µm cell strainer. The cell strainer was washed twice and the cell suspension was centrifuged at 500g for 5 min before supernantant was removed and the pellet was frozen in 1 ml Cryobrew (cat. no.: 130-109-558, StemMACS, Miltenyi Biotec) before further analysis. Biopsies were obtained pre-injury as well as 2h, 2, 8, and 30 days after injury. FAP quantification from one individual from the injury study was excluded from further analysis due to exceptional high FAP count in the pre sample.

Skeletal muscle tissue for *ex vivo* studies (C3 ELISA and C3 staining of isolated cells) was obtained from donors undergoing leg amputation at Aarhus University Hospital, Aarhus, Denmark. The biopsies were obtained from the amputated part as close to the vital amputation line as possible.

Monocytes from peripheral blood for *ex vivo* studies of monocytes and macrophages were obtained from three healthy male donors.

### Spatial transcriptomics and bioinformatics

Spatial transcriptomics analysis of skeletal muscle was carried out on samples from three subjects (gender: female; age: 57-59; BMI: 23-28 kg/m2; no chronic diseases; received no medication). Biopsies from pre, 2, 8, and 30 days post injury were embedded with OCT and 10 µm thick cryosections were used for analysis. The cryosections were placed directly on Visium slides (cat. no.: PN-1000184, 10X Genomics) with one slide containing pre, 2, 8, and 30 days post injury samples from the same donor. The spatial sequencing libraries were prepared following the manufacturer’s instructions of Visium Spatial Gene expression reagent kits guide (CG000239, 10X Genomics). After library construction, the library conversion was performed using the MGIEasy Universal DNA Library Preparation reagent kit (BGI, Shenzhen, China) for compatibility, followed by sequencing on a DNBSEQ-G400 platform (MGI).

#### Preprocessing and quality control of spatial-sequencing data

Space Ranger1.3.0 (10X Genomics) was used to process the raw sequencing data. The detailed report of sequencing and alignment is available through our data-base website. To ensure the quality of data, the generated gene expression matrix was filtered if the UMI count within one spot was less than 700.

#### Identification of spot clusters and cluster markers

After filtering, unsupervised clustering was performed using Seurat (v4.0.3, Satija et al., 2015). Briefly, datasets from different sequencing libraries were merged after SCT transformation. After principal components analysis, clusters were identified via the FindClusters function at resolution 1 with PC1-PC30 in Seurat and subsequently visualized using the RunUMAP function (reduction = “pca”). “FindAllMarker” function implemented in Seurat was used to identify DEGs across clusters with the options “min.pct = 0.25, logfc.threshold = 0.25”. Adjust P value of 0.05 was set as a threshold to define significance. Furthermore, cluster annotation was assigned by using known region markers.

#### Gene ontology (GO) enrichment analysis

The gene lists were subjected to Gene Ontology analysis using the enrichGO function implemented in the clusterProfiler package (v3.18.1). Gene Ontology enrichment level was evaluated by adjusted P values, and multiple test adjustment was conducted using the Benjamini-Hochberg method.

#### Single-cell RNA sequencing data preparation for decomposition

Briefly, single-cell data from a previously published study (30) were cleaned up by removing debris (gene count <200), doublet (gene count > 4000), and dying cells (mitochondrial gene percentage > 15%). Afterward normalization, log1P transformation, and top 2000 variable gene detection, data from different samples were integrated with FindIntegrationAnchors and IntegrateData function in Seurat. After principal component analysis and identification of clusters, scRNAseq data were annotated based on the previous reported markers.

#### Decomposition and cell type correlations

Each of the spots were decomposed with the SPOTlight package (v0.1.7). Annotated scRNAseq data were SCT transformed, followed by principal component analysis and dimension deduction with UMAP. Cell type markers were identified with FindAllMarkers function in Seurat and used for decomposition of spatial data with spotlight_deconvolution function using the parameter of (cl_n = 100, hvg = 3000, ntop = NULL, transf = “uv”, method = “nsNMF”, min_cont = 0). Pearson analysis was used to analysis the cell type correlation in the spot, and p-values and confidence were calculated with a significance test using cor.mtest function in corrplot package (v0.89).

#### Cell-Cell communication analysis

Cell-cell communication analysis was performed with CellChat (v1.1.3) with scRNAseq data. Briefly, communication probability and intercellular communication network were computed with omputeCommunProb function. Cell-cell communication was inferred at a signaling pathway level using computeCommunProbPathway function and aggregated with aggregateNet function.

#### Data availability

Processed data are available and visualized in DREAMapp (https://dream.au.dk/tools-and-resources/humdb).

### Immunohistochemistry

10 µm thick cryosections from pre, 2, 8, and 30 days post injury were used for analysis. Sections were thawed to room temperature and added Histofix (Histolab Products AB, Västra Frölunda, Sweeden) for 5 min. Histofix was then removed and sections were washed once in 1X PBS followed by 1% BSA + 10% FBS + 0.5% Triton in 1X PBS for 60 min at room temperature for blockade of unspecific binding and permeabilization. Sections were incubated overnight with specific antibodies targeting CD11c (rabbit anti human, 1:100, cat. no.: ab52632), CD68 (mouse anti human, 1:100, cat. no.: M0718, Dako Norden, Glostrup Denmark), PDGFR-α (goat anti human, 1:300, cat. no.: AF-307-NA; RRID: AB_354459, R&D Systems, Bio-Techne), and C3 (rabbit anti human, 1:100, cat. no.: PA5-21349, Thermo Fisher Scientific). Thereafter, sections were washed once in 1X PBS and incubated with Wheat Germ Agglutinin (WGA) Texas Red™-X Conjugate (1:500, cat. no.: W21405, Thermo Fisher Scientific) and secondary antibodies (Goat-anti-mouse or Mouse-anti-rabbit Alexa Flour 488, 587 or 647) in 1X PBS + 1% BSA for 60 min at room temperature and washed in PBS 3×5 min, with one wash containing DAPI (1:50.000, cat. no.: D3571, Invitrogen, Thermo Fisher Scientific). Minus primary controls were included for all stains during optimization to ensure specificity. Finally, the section was mounted with cover slides. EVOS M7000 automated imaging system (Thermo Fisher Scientific) was used for image capturing.

### Cell isolation and fluorescence activated cell sorting

Samples were thawed in a 37 deg C water bath and added 10 ml of wash buffer before centrifuged at 500g for 5 min. The supernatant was removed, and the pellet was resuspended in 400 µl wash buffer and added:

- FcR-block (20µl/sample, cat. no.: 130-059-901, Miltenyi Biotec)
- Anti-CD56-BV421 (5µl/sample, cat. no.: 562751, BD Biosciences, CA, USA)
- Anti-CD82-PE-Vio770) (10µl/sample, cat. no.: 130-101-302, Miltenyi Biotec)
- Anti-CD34-APC (20µl/sample, cat 555824, BD Biosciences)
- Anti-CD31-PerCPvio 700 (2µl/sample, cat. no.: 700 130-110-673, Miltenyi Biotec)
- Anti-CD90-PE (3,6µl/sample, cat. no.: 12-0909-42, Invitrogen, ThermoFisher Scientific)
- Anti-CD45-VioBright FITC (12µl/sample, cat. no.: 130-114-567, Miltenyi Biotec)
- Anti-CD14-BV605 (5µl/sample, cat. no.: 564054, BD Biosciences)
- Anti-CD11c-APCvio770 (2µl/sample, cat. no.: 130-113-585, Miltenyi Biotec)

The suspension was incubated at 5 deg C for 30 min protected from light before washed in 10 ml wash buffer and centrifuged at 500g for 5 min. The supernatant was removed, and the pellet was resuspended in wash buffer and filtered through a 30 µm cell strainer. For viability staining, propidium iodide (PI, 10µl/sample, cat. no.: 556463, BD Bioscience) was added before flow cytometry. For absolute quantification, CountBright™ counting beads (10µl/sample, cat. no.: C36950, ThermoFisher Scientific) were used together with total milligram of tissue.

MuSCs were quantified and sorted as CD31-CD45-CD34-CD56+CD82+, FAP as CD31-CD45-CD34+, monocytes as CD45+CD14+CD11c+/- and endothelial cells as CD45-CD31+. Lymphocytes were defined as CD45+ with small size and granularity (Forward-scatter low, Side-scatter low).

Primary human monocytes were isolated from healthy human blood samples using CD14-magnetic beads (cat. no.: 130-050-201, Miltenyi Biotec). In brief, following sample collection, the erythrocytes were lysed in a lysis buffer (cat. no.: 130-094-183, Miltenyi Biotec) for 1 min. Leukocytes were then centrifuged and washed once in PBS. Following this the leukocytes were incubated with anti-CD14-Microbeads or anti-CD14-BV421 (5ul/test, Cat. No.: 564943, BD) in stain buffer (PBS + 2mM EDTA + 1% BSA) for 15 min at room temperature. The cells were then washed twice in stain buffer and applied to a magnetic column (cat. no.: 130-042-201, Miltenyi Biotec). The column was washed x 3 with stain buffer, removed from the magnet, and column retained cells were flushed out. Flow-cytometry analysis of flow-through (CD14-) and the retained cells (CD14+) revealed a marked enrichment for CD14+ monocytes (**Supplementary Figure 6A**). For phagocytosis experiments enriched monocytes were plated in ECM-coated wells (54000/cm2) in Advanced RMPI 1640 medium (cat. no.: 12633012, Gibco, Thermo Fisher Scientific) + 10% FBS (cat. no.: 16000044, Thermo Fisher Scientific) + 50 ng/ml M-CSF (cat. no.: 130-096-491, Miltenyi Biotec) + 1% Penicillin-Streptomycin (cat. no.: 15140122, Gibco, ThermoFisher Scientific, MA, USA) for six days to induce differentiation into macrophages.

### ELISA

For C3-ELISA experiments, FAPs, endothelial cells and MuSCs were isolated from human skeletal muscle and plated as 200 cells/µl medium (HAMS F10 + 5% FBS + 1% Penicillin-Streptomycin (cat. no.: 15140122, Gibco, ThermoFisher Scientific, MA, USA) and incubated for 48 h. The medium was then collected and stored at - 80°C until further analysis. Cells were fixed in 4% paraformaldehyde for 8 min followed by 3x 5 min 1X PBS before storage at 5°C. Cell media was analyzed with ELISA for C3 protein as prescribed by manufacturer (cat. no.: CS409A, Cell Sciences, US). Fixed cells were stained for C3 protein using the same procedure as for immunohistochemistry.

### Phagocytosis assay

Following six days in macrophage induction medium, monocytes uniformly expressed CD68 (**Supplementary Figure 6B**). For phagocyte assay, C2C12 myoblasts were labelled with a lipophilic dye for 2 minutes at room temperature (cat. no.: PKH67, Sigma, MINI67) as per manufacturer instructions. C2C12’s were replated and allowed 24 hours to recover after which the labeled C2C12 myoblasts were induced for apoptosis using 5 µM Staurosporin (cat. no.: 9953S, Cell Signaling) for 18h. Hereafter they were added to macrophages stimulated with or without C3 (cat. no.: CAS 80295-41-6, Merck KGaA, Germany) in a macrophage:C2C12 ratio of 1:3 for 4 hours at 37 deg C. A separate plate was incubated at 4 deg C as a negative control. Following phagocytosis, the cells were washed 6 times in PBS and fixed/permeabilized. To identify macrophages the cells were incubated with anti-CD68 BV421 (??) for 30 min at 4 deg, after which the cells were analyzed on a Penteon Analyser (Agilent). Phagocytosis level was determined by PKH67 MFI expression in CD68-PKH67 positive cells.

### Cell proliferation

FAP proliferation was detected using 5-ethynyl-2’-deoxyuridine (EdU)/click-it assay (cat. no.: C10337 and C10340, Invitrogen). Experiments were carried out in ECM coated 96-well half-area plates. FAPs were plated in wash buffer+10µM EdU immediately after sorting. After 48h, cells were fixed in 4% paraformaldehyde for 8min, washed 3×5min in PBS and kept in PBS at 4°C. To detect EdU-incorporation, we followed manufacturer protocol and counter-stained with DAPI (1:50.000, D3571, Invitrogen, Thermo Fisher Scientific). Images were acquired using an EVOS M7000 automated imaging system (Thermo Fisher Scientific). EdU- and DAPI-positive cells were semi-automatically counted in ImageJ.

### Cell migration

Migration of directly isolated CD14^+^ cells was carried out using ECM coated µ-Slide Chemotaxis (cat. no.: 80326, Ibidi GmbH, Germany). 6 µl of cell mixture (RPMI + 0.5% FBS + 1×10^6^ directly isolated CD14+ cells) was added to the chamber followed by RPMI + 0.5% FBS in the reservoirs. C3 in 1xPBS were then applied to the reservoirs as either 50 µg/mL, 500 µg/mL or 1xPBS in an equal volume as control. Hereafter, time lapse microscopy was conducted on EVOS M7000 automated imaging system (Thermo Fisher Scientific) with an on-stage incubator for 12h, one image per 3 min. 30 cells per intervention were manually tracked using ImageJ.

### Cell survival

For survival experiments, CD14^+^ cells were directly plated in ECM-coated 12-wells in Advanced RMPI 1640 medium (12633012, Gibco, Thermo Fisher Scientific) + 1% Penicillin-Streptomycin (cat. no.: 15140122, Gibco, ThermoFisher Scientific, MA, USA) + 10 % FBS (high serum) or 1% FBS (low serum) ± C3 (10 or 100 µg/ml) for seven days. Cells were then stained directly in wells with Hoechst (cat. no.: H3570, Invitrogen, ThermoFisher Scientific, MA, USA) and Propidium Iodide (cat. no.: P2667, Merck KGaA, Germany) in 1X PBS for 20 min followed by imaging. Cells positive for Propidium Iodide and/or Hoechst staining were semi-automatically quantified using ImageJ.

### Bioenergetic analysis

Realtime analysis of oxygen consumption rate (OCR) and extracellular acidification rate (ECAR), as a proxy of cellular glycolysis, was performed using the Seahorse technology.

Monocytes were isolated and differentiated into macrophages as described. The cells were exposed to either C3 (or control) or apoptotic C2C12 for 6 hours followed by extensive washing. For bioenergetic analysis, we plated 1×10^5^ macrophages in ECM (Sigma) coated Seahorse XF HS Mini cell culture microplate (part. no.: 102984-000, Agilent) in wash-buffer (to minimize change in cell bioenergetics related to high concentration of growth-factors etc). The number of cells was optimized before running the experiment. One hour before the analysis, the cells were washed twice in phenol red and bicarbonate free DMEM Seahorse Media (SM) (part. no.: 103575-100, Agilent) containing 25 mM glucose (Cat no. G8769, Sigma), 1 mM sodium-pyruvate (cat no.: 11360-039, Gibco) and 2 mM L-glutamine (cat no.: G7513, Sigma), (pH 7.4) and allowed to equilibrate in a non-Co2 incubator for an hour. During this time brightfield images of all wells were captured to ensure a sufficient monolayer of cells. For directly isolated cells, we performed the bioenergetic analysis using the Seahorse T-cell Mitochondrial profiling kit (part. no.: 103771-100, Agilent) with a final concentration of 4mM Oligomycin, 2mM BAM15 and 2.5 mM Antimycin A/Rotenone diluted into the SM media. Bioenergetic analysis was performed on the Seahorse XF HS Mini Analyzer (Agilent). Following the bioenergetic analysis Hoechst 33342 (cat. no.: H3570, Invitrogen) was added to all wells and images obtained of all wells to count the number of cells in each well. The number of cells per well was utilized to normalize OCR and ECAR values.

### Statistics

Data are expressed as mean ± standard deviation if normally distributed or geometric mean ± standard deviation if not normal distributed. Data distribution was evaluated by inspection of QQ-plots. Absolute cell quantification by FACS (FAPs, total monocytes, lymphocytes, MuSCs, endothelial cells and CD11c^+/-^- monocytes after log-transformation), phagocytosis, cell death, cell size and EdU-data were analyzed using a one-way ANOVA. Bioenergetic data were analyzed using a paired t-test. Significance was set at p<0.05. All analyses were carried out using GraphPad Prism version 8.3.0 (GraphPad Software) and STATA version 16. Graphs were generated in GraphPad Prism and figures were created with Biorender.com. Specific analysis for transcriptomic data are stated in Spatial transcriptomics and bioinformatics.

## Notes

https://dream.au.dk/tools-and-resources/humdb

### Competing Interest Statement

The authors have declared no competing interest.

