## Supplementary data for "Complementing Muscle Regeneration: Fibro-Adipogenic Progenitor and Macrophage-Mediated Repair of Elderly Human Skeletal Muscle"

A

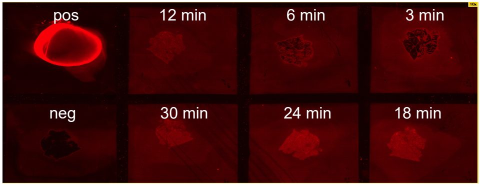


B

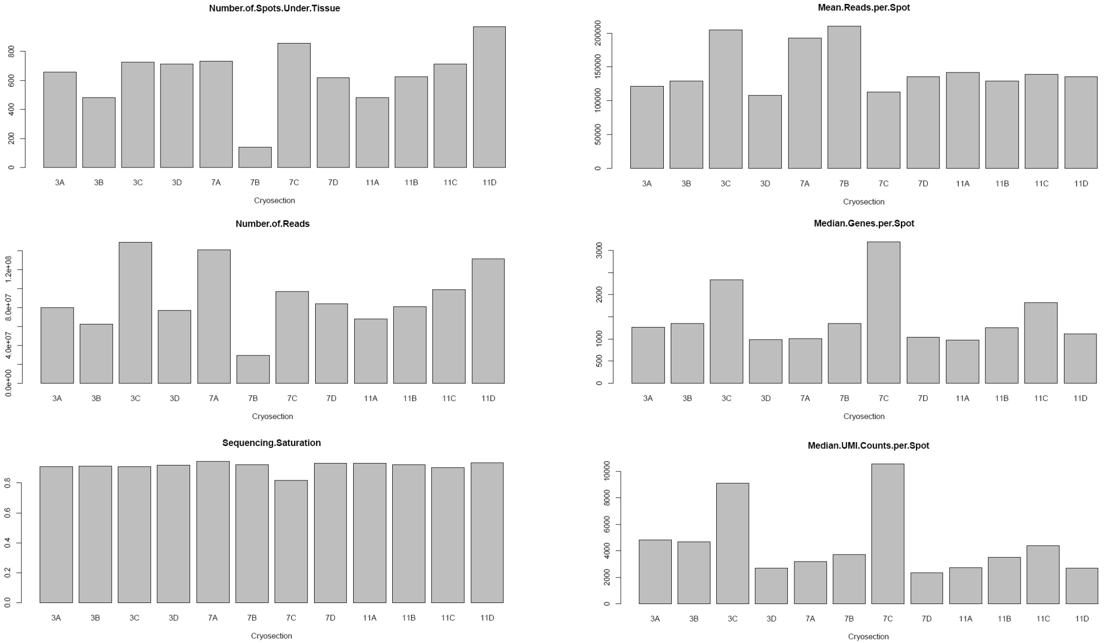


**Supplementary Figure 1:**
A: cDNA staining of skeletal muscle cryosections. Time points represent permabilization time. B: Quality control of spatial sequencing data. pos and neg = positive and negative control, respectively.

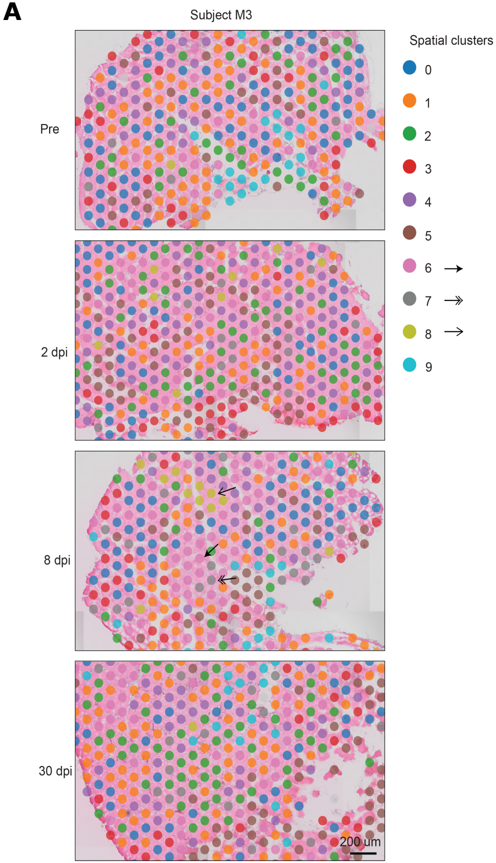


**Supplementary Figure 2:**
A: Spatial spots divided into nine different clusters based on gene expression illustrating spots contaning cluster 6, 7, and 8 are more present after injury.

**A**

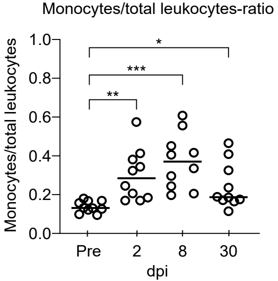


**Supplementary Figure 3:**

A: Monocytes/total leukocytes-ratio. * = $p < 0.05$. ** = $p < 0.01$. *** = $p < 0.001$. dpi = days post i

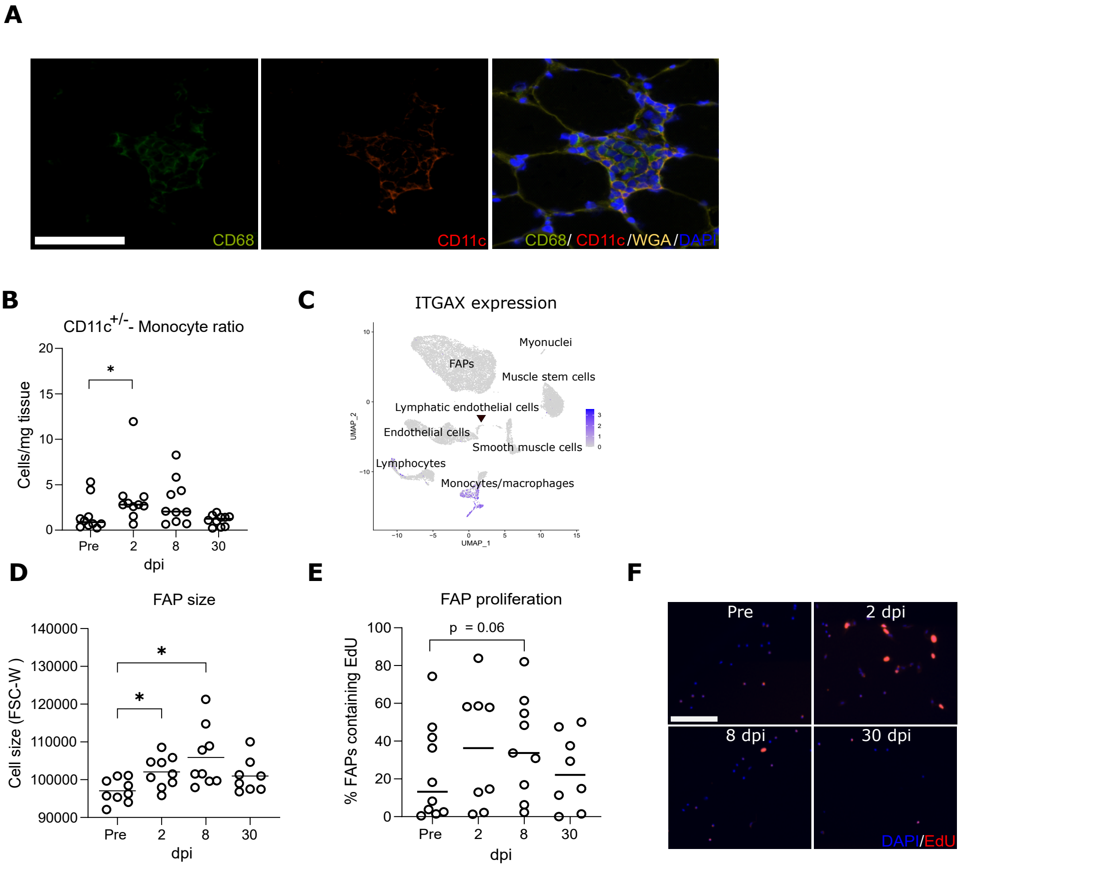


**Supplementary Figure 4:**

A: Immunostaining of CD68, CD11c, WGA, and DAPI 8 dpi. Scale bar represents 125 µm. B: CD11c$^{+/-}$-monocyte ratio (n = 10). C: tSNE-plot of ITGAX expression in single cells. D: FAP size determined by flow cytometry (n = 9). E+F: EdU-positive FAPs 48h after isolation (n = 9). Scale bar represents 275 µm. * = $p < 0.05$. dpi = days post injury.

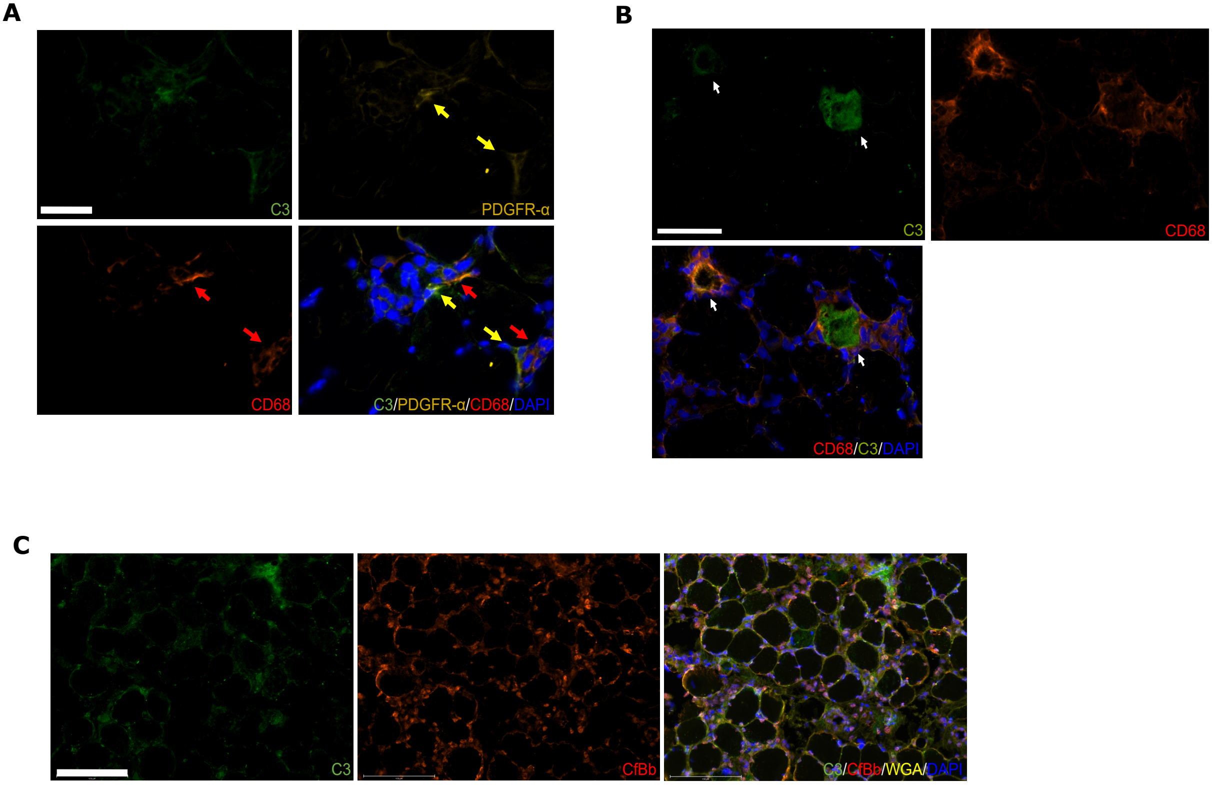


**Supplementary Figure 5:**
A: Immunostaining of C3, PDGFR-α, CD68, and DAPI 8 dpi. Scale bar represents 75 µm. B: Immunostaining of C3, CD68, and DAPI 8dpi. Scale bar represents 125 µm. C: Immunostaining of C3, CfBb, WGA, and DAPI 8 dpi. Scale bar represents 150 µm.

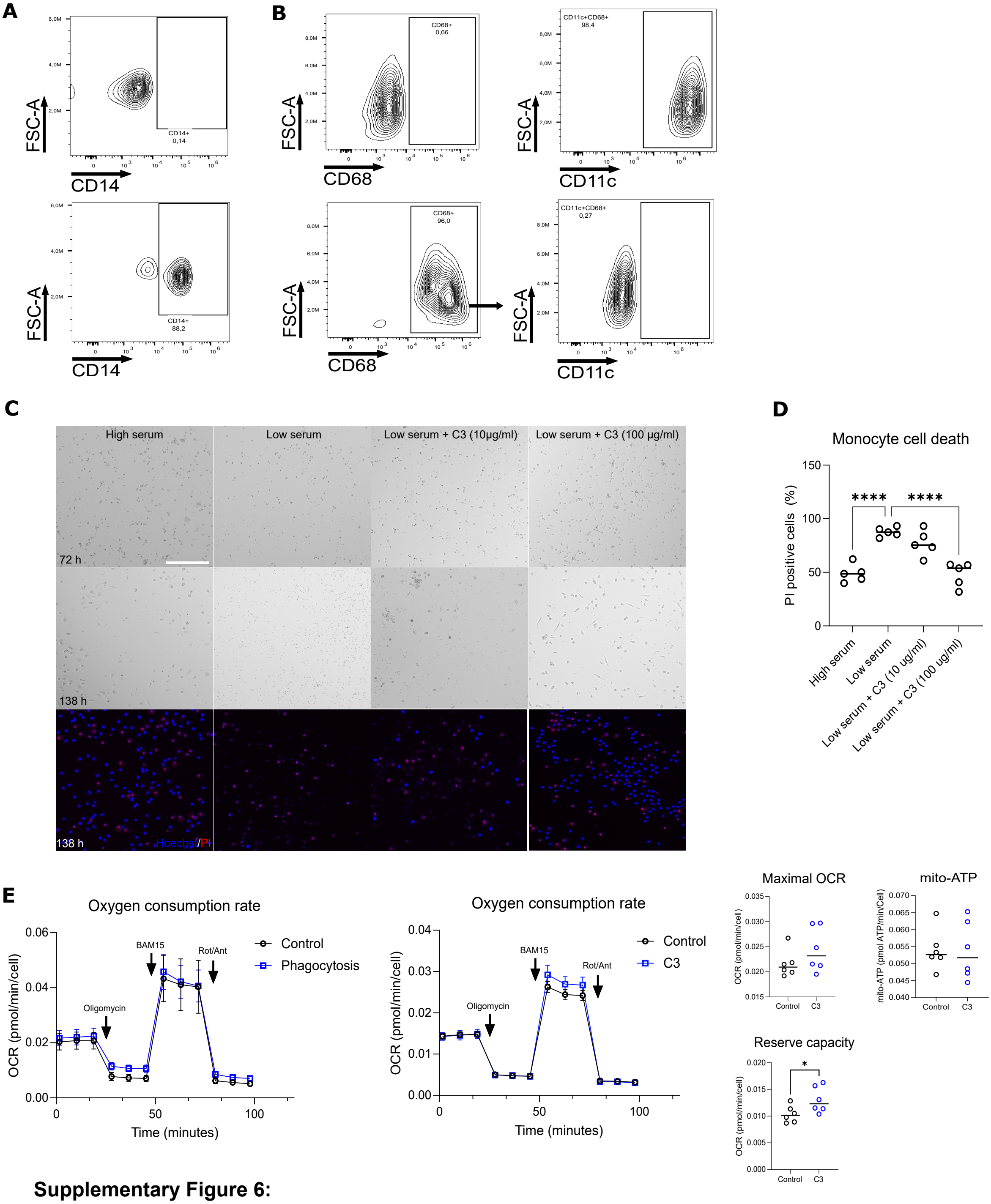


**Supplementary Figure 6:**

A: Flow plots of directly isolated monocytes from human peripheral blood with unstained control (above) and stained cells (below). B: Flow plots of isolated monocytes from human peripheral blood after 6 days of stimulation with monocyte colony stimulating factor, unstained control (above) and stained cells (below). Flow plots of isolated monocytes from human peripheral blood after 6 days of stimulation with monocyte colony stimulating factor confirming the majority is positive for CD11c , unstained control (ab ove) and stained cells (below). C+D: Cell death of directly isolated monocytes in response to low serum revealing an increased survival with C3 protein and a markedly different morphology . **** = p < 0.0001. Scale bar represents 275 µm. E: Measurement of oxygen consumption rate (OCR), mitochondrial ATP and reserve capacity in macrophages stimulated with C3 protein alone or undergoing C3 mediated phagocytosis.

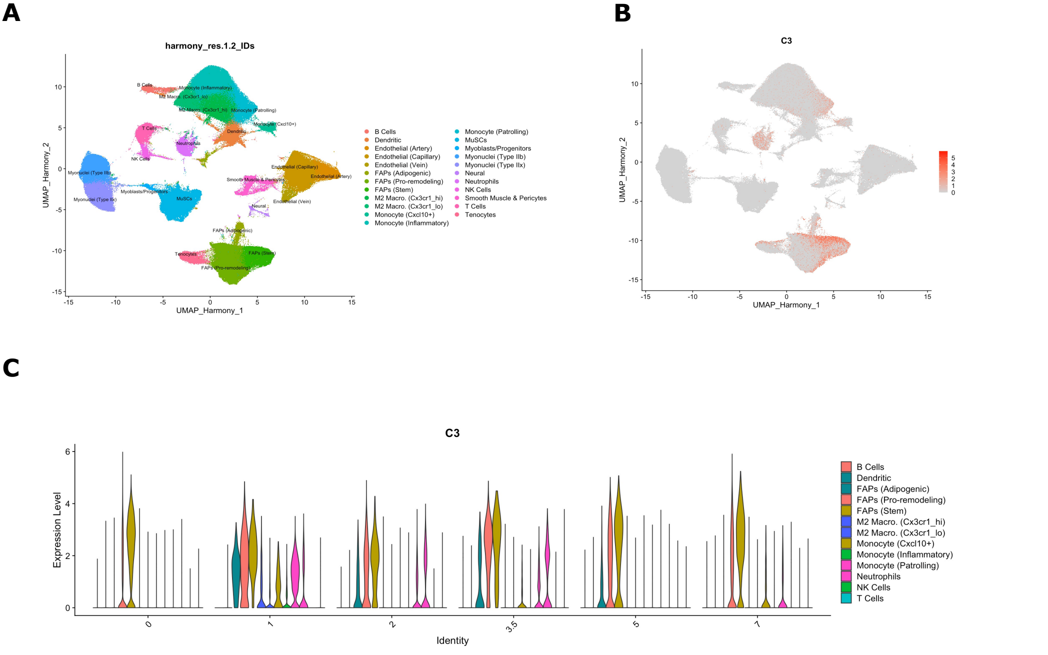


**Supplementary Figure 7:**
A: UMAP plot of murine single-cell sequencing data from mononucelar cells in skeletal mucle. B: UMAP plot of C3 expression in single cells. C: C3 expression in FAPs and immune cells in murine skeletal mucle injury and regeneration.
